# IgG hexamers initiate acute lung injury

**DOI:** 10.1101/2024.01.24.577129

**Authors:** Simon J. Cleary, Yurim Seo, Jennifer J. Tian, Nicholas Kwaan, David P. Bulkley, Arthur E. H. Bentlage, Gestur Vidarsson, Éric Boilard, Rolf Spirig, James C. Zimring, Mark R. Looney

**Affiliations:** Department of Medicine, University of California, San Francisco (UCSF), CA, USA; Department of Biochemistry and Biophysics, University of California, San Francisco (UCSF), CA, USA; Sanquin Research, Amsterdam, The Netherlands; Centre de Recherche du Centre Hospitalier Universitaire de Québec - Université Laval, Québec, QC, Canada; CSL Behring, Research, CSL Behring Biologics Research Center, Bern, Switzerland; Department of Pathology, University of Virginia School of Medicine, Charlottesville, VA, USA

## Abstract

Antibodies can initiate lung injury in a variety of disease states such as autoimmunity, transfusion reactions, or after organ transplantation, but the key factors determining in vivo pathogenicity of injury-inducing antibodies are unclear. A previously overlooked step in complement activation by IgG antibodies has been elucidated involving interactions between IgG Fc domains that enable assembly of IgG hexamers, which can optimally activate the complement cascade. Here, we tested the in vivo relevance of IgG hexamers in a complement-dependent alloantibody model of acute lung injury. We used three approaches to block alloantibody hexamerization (antibody carbamylation, the K439E Fc mutation, or treatment with domain B from Staphylococcal protein A), all of which reduced acute lung injury. Conversely, Fc mutations promoting spontaneous hexamerization made a harmful alloantibody into a more potent inducer of acute lung injury and rendered an innocuous alloantibody pathogenic. Treatment with a recombinant Fc hexamer ‘decoy’ therapeutic protected mice from lung injury, including in a model with transgenic human FCGR2A expression that exacerbated pathology. These results indicate a direct in vivo role of IgG hexamerization in initiating acute lung injury and the potential for therapeutics that inhibit or mimic hexamerization to treat antibody-mediated diseases.

**Brief summary:** IgG antibodies can form hexamers. This study shows that hexamer assembly is an important event determining the ability of IgG to trigger acute lung injury.

**Graphical abstract:** 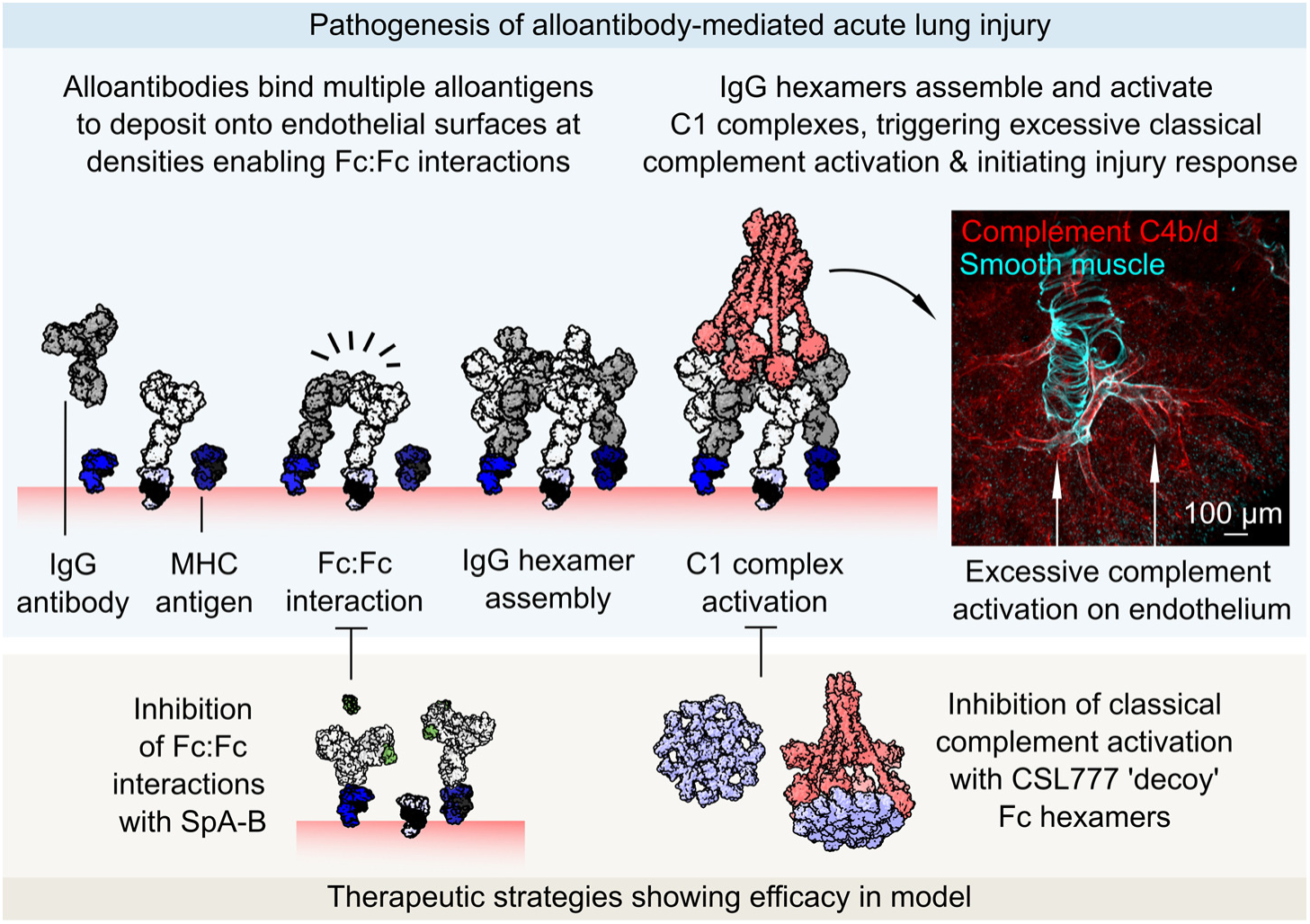

## Introduction

Antibodies and the complement cascade mediate protective immunity but can both become misdirected to cause harm in autoimmune and alloimmune diseases. Some antibodies can direct activation of the complement cascade at their targets, an event associated with severe pathology in disease states including several forms of transfusion reactions (1, 2), immune rejection after solid organ transplantation (3), and complications of pregnancy (4). Complement-activating alloantibodies are known mediators of transfusion-related acute lung injury (TRALI) (5, 6), a leading cause of transfusion-related deaths (7), and are linked to particularly poor outcomes following solid organ transplantation (3, 8). Complement activation by autoreactive antibodies also contributes to pathogenesis of forms of autoimmune hemolytic anemia (9), small vessel vasculitis (10), and neurological autoimmune disease (11).

Immunoglobulin G (IgG) antibodies are the most prevalent type of antibody in circulation and complement-activating alloantibodies are frequently IgG class. IgG antibodies achieve complement activation through recruiting and activating C1 complexes, each of which contain six Fc-binding domains (12, 13). A theory for how IgG achieves C1 complex activation involves groups of six IgG antibodies interacting through their Fc domains to form IgG hexamers (14). This theory recently gained experimental support from direct imaging of IgG1 and IgG3 hexamer assembly on antigenic liposomes (15, 16), with in vitro studies connecting IgG hexamerization to increased complement deposition on target surfaces (15, 17, 18). However, it is currently unclear whether IgG hexamer assembly is important in vivo in the pathogenesis of complement-dependent forms of alloantibody-mediated disease.

Here, we report testing of interventions that exploit IgG hexamerization in a mouse model of acute lung injury driven by alloantibody deposition in the pulmonary microvasculature, a process that drives pathology in forms of both TRALI and antibody-mediated rejection (AbMR) of lung transplants (6). Our results identify key molecular events driving alloantibody-mediated pathophysiology in vivo. We also demonstrate preclinical efficacy of new therapeutic approaches that prevent pathology of complement-dependent organ damage caused by alloantibodies, serving as a rationale to pursue translational studies in human alloantibody driven disease.

## Results

Alloantibodies are prevalent but not always harmful, so determining whether alloantibodies are clinically significant is a frequent conundrum in transfusion and transplantation medicine. Reflecting this clinical challenge, of the many mouse monoclonal alloantibodies targeting major histocompatibility complex (MHC) class I antigens, only clone 34-1-2S triggers acute lung injury when microgram quantities are intravenously injected into mice (19, 20). In addition, only mice expressing the H-2^d^ MHC class I haplotype are known to be susceptible to injury caused by the 34-1-2S antibody (5, 6). Curiously, the 34-1-2S antibody does not readily cause injury in H-2^b^ mice, including the widely used C57BL/6 (B6) strain, despite the fact that it binds to MHC class I antigens expressed by H-2^b^ mice (5, 6). We aimed to measure the affinity of 34-1-2S for a range of MHC class I antigens to improve our understanding of the factors determining the ability of antibodies to cause injury in both this widely used model and more generally in antibody-mediated disease states.

We measured the binding affinity of 34-1-2S antibody to each of the classical MHC class I antigens present on injury-resistant H-2^b^ B6 mice and injury-susceptible H-2^d^ mice (**Fig. 1A**). Of the three MHC antigens in the H-2 locus (K, D, and L), B6 mice only express K^b^ and D^b^, and we detected binding of 34-1-2S to K^b^ but not D^b^. In contrast, we detected binding of 34-1-2S to all three MHC class I antigens from H-2^d^ mice, with high affinity binding to K^d^ and D^d^, and weak binding to L^d^ (**Fig. 1B**). Other MHC class I antibodies (clones AF6-88.5.5.3, 20-8-4S, SF1.1.10, 30-5-7S and 34-5-8S), which do not readily induce injury (5)) each bound to only one MHC class I antigen from each MHC type (**Fig. S1**).

**Fig. 1.**
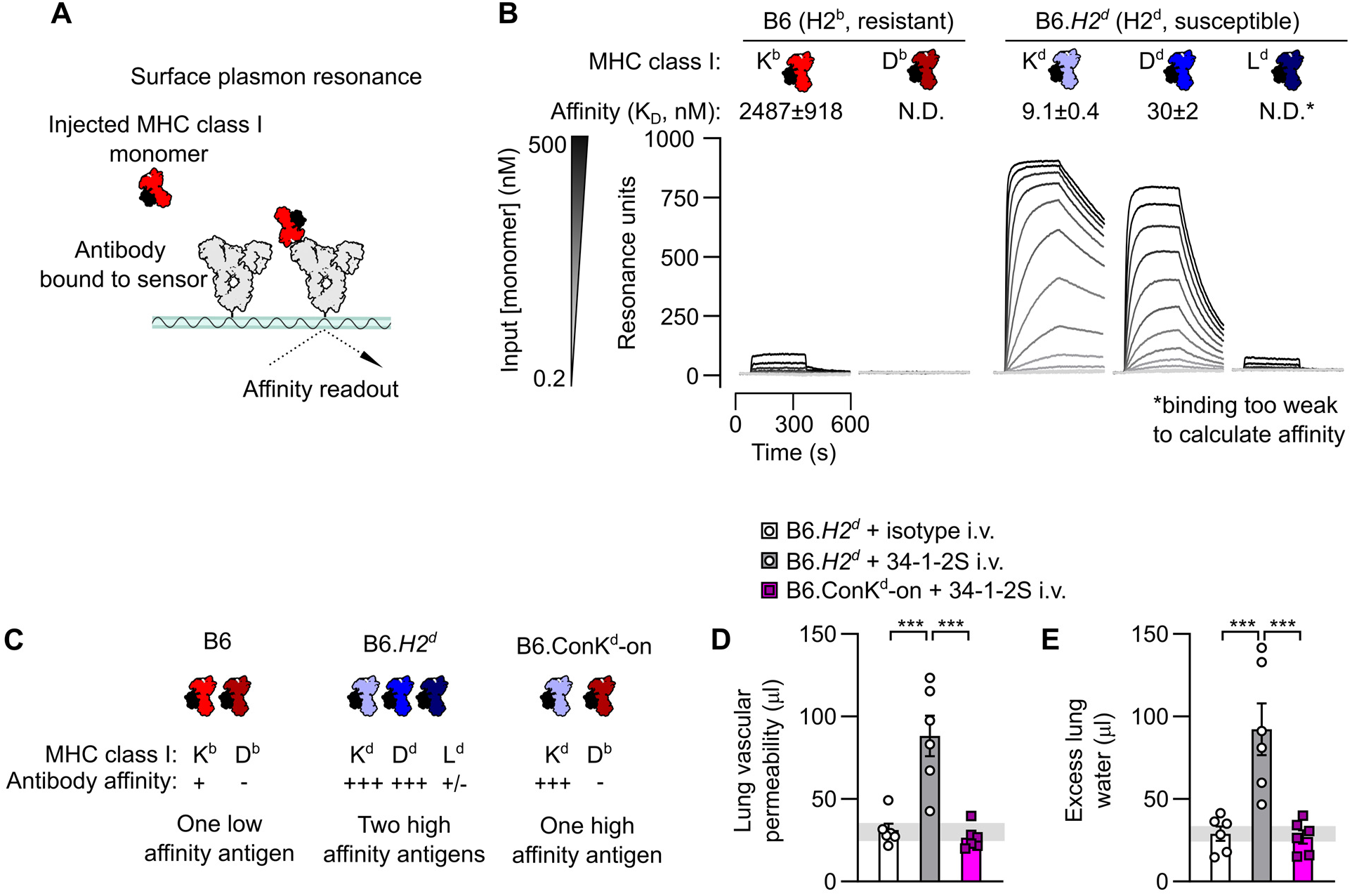
The 34-1-2S alloantibody binds to multiple MHC class I antigens to trigger acute lung injury. **A.** Schematic showing approach for measuring affinity of the MHC class I alloantibody 34-1-2S for MHC class I monomers using surface plasmon resonance (SPR). **B.** SPR sensorgrams showing detection of binding of 34-1-2S to the K^b^ MHC class I antigen from H2^b^ mice and the K^d^, D^d^ and L^d^ antigens from mice with the H2^d^ haplotype. **C.** Classical MHC class I antigens present in B6, B6.*H2^d^* and B6.Con-K^d^-on mice with summary of results from **B**. **D.** Lung vascular permeability and **E**. excess lung water measurements from LPS-primed B6.*H2^d^* mice given intravenous (i.v.) doses of either 34-1-2S or isotype control, versus B6.Con-K^d^-on mice given i.v. 34-1-2S. Depictions of IgG and MHC class I in **A-C** are based on protein data bank (PDB) entries 1HZH and 1RK1. **B, D & E** show means ± standard errors. Statistical tests used on **D** & **F** were ordinary one-way ANOVA with Dunnett’s test for differences relative to B6.*H2^d^* + 34-1-2S i.v. group, with data log_10_-transformed prior to analysis, *** = *P*<0.0001.

Together, the above findings led us to the hypothesis that the ability of 34-1-2S to induce lung injury in H-2^d^ mice is a function of increased density of bound antibody in H-2^d^ animals resulting from 34-1-2S simultaneously binding K^d^, D^d^, and possibly L^d^. This hypothesis was tested by injecting 34-1-2S antibody into B6.ConK^d^-on mice, which express K^d^ but do not express D^d^ or L^d^ (21). B6.*H2^d^* mice expressing the full complement of MHC class I antigens recognized by 34-1-2S (K^d^, D^d^, and L^d^) were used as background-matched positive controls for susceptibility to injury (**Fig. 1C**). In contrast to B6.*H2^d^* mice, B6.ConK^d^-on mice did not develop lung injury (**Fig. 1D, E**). These data are consistent with 34-1-2S antibody causing injury in H2^d^ mice through high affinity binding to multiple MHC class I antigens.

Engagement of multiple antigens can permit high density antibody deposition, an event associated with classical complement activation. Complement activation has been implicated in pathogenesis of acute lung injury caused by 34-1-2S antibody, but previous studies have not determined whether injury in this model is directly triggered by antibody-mediated complement activation via the classical pathway (5, 6).

To test whether 34-1-2S-induced injury requires classical complement activation, we bred mice expressing the *H2^d^* susceptibility locus with mice lacking C1qa (22), a protein that is necessary for classical complement activation as it is one of the three proteins which make up each of the six Fc-binding C1q subcomponents in each C1 complex (**Fig. 2A**).

**Fig. 2.**
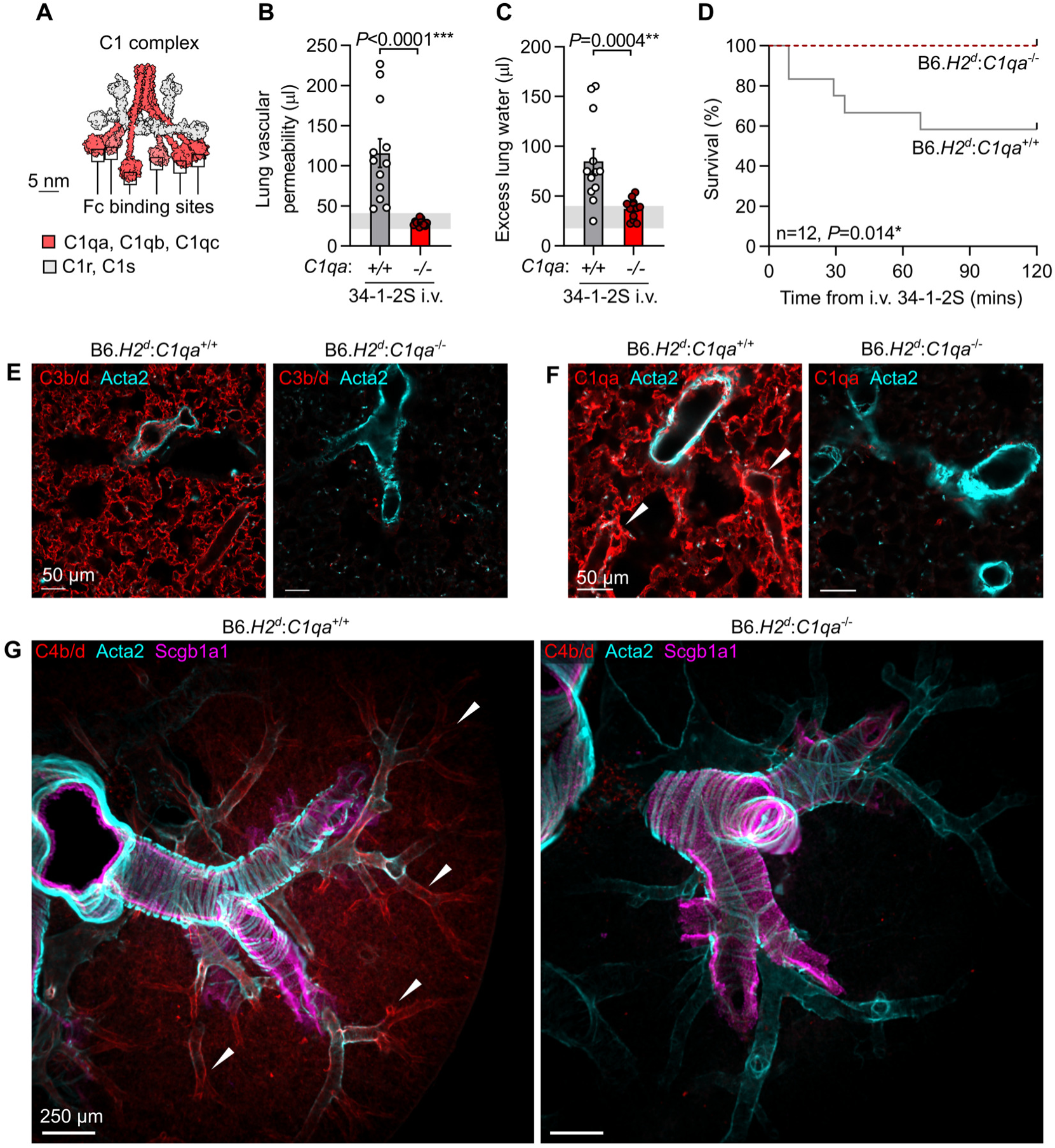
Classical complement activation on the pulmonary endothelium initiates 34-1-2S-induced acute lung injury. **A.** Molecular model of C1 complex based on small angle scattering database entry SASDB38 (12). **B.** Lung vascular permeability and **C**. excess lung water measurements from LPS-primed B6.*H2^d^*:*C1qa*^-/-^ mice and B6.*H2^d^*:*C1qa*^+/+^ littermates given i.v. 34-1-2S at 1 mg/kg. Horizontal gray lines are standard deviations of values from ‘no injury’ controls (B6.*H2^d^* mice given LPS i.p. + mIgG2a isotype control i.v.) **C.** Survival of LPS-primed B6.*H2^d^*:*C1qa*^-/-^ mice and B6.*H2^d^*:*C1qa*^+/+^ littermates given i.v. 34-1-2S at 4.5 mg/kg. **D.** Immunofluorescence staining for complement C3b/d, **F**. C1qa or **G**. C4b/d (red) as well as Acta2 (α-smooth muscle actin, cyan) and, in **G**., Scgb1a1 (club cell secretory protein, magenta) in lung sections from LPS-primed B6.*H2^d^*:*C1qa*^-/-^ mice and B6.*H2^d^*:*C1qa*^+/+^ mice fixed 5 minutes after i.v. 34-1-2S at 1 mg/kg. Images in **G**. are maximum intensity projections sampling 240 µm from a cleared thick section. White arrowheads point to arterioles positive for complement components. **B** & **C** show means ± standard errors. *P*-values are from unpaired two-tailed t-tests on log_10_-transformed data (**B** & **C**) or log-rank test (**D**), with group n=12.

Relative to B6.*H2^d^*:*C1qa*^+/+^ littermate controls, C1qa-deficient B6.*H2^d^*:*C1qa*^-/-^ mice were resistant to alloantibody-mediated acute lung injury and mortality (**Fig. 2B-D**). Mice lacking C1qa were also protected from deposition of complement component C3 split products on the endothelium of pulmonary capillaries (**Fig. 2E**). Staining for C1qa in lungs confirmed absence of C1qa protein in knockout mice, with intense C1 complex deposition seen around pulmonary arteriolar endothelial cells in *C1qa*-expressing mice injected with 34-1-2S antibody (**Fig. 2F**).

To identify the microanatomical site of classical complement activation, we stained lungs of mice injected with 34-1-2S for the complement split products C4b and C4d, which form covalent bonds with proteins at sites of classical complement activation. We observed strong positivity for C4b/d highlighting the endothelium of medium and small-sized pulmonary arterioles in B6.*H2^d^*:*C1qa*^+/+^ mice injected with 34-1-2S, but not in B6.*H2^d^*:*C1qa*^-/-^ mice (**Fig. 2G** and **Movie S1**). Together, these results indicate that 34-1-2S causes acute lung injury because this antibody is deposited onto the pulmonary arteriolar endothelium at densities sufficient to trigger excessive classical complement activation directed at the walls of these blood vessels.

Dense binding to membrane-expressed antigens would be expected to facilitate IgG Fc:Fc interactions and IgG hexamer assembly. IgG hexamers are potent activators of C1 complexes in vitro (15), and are further implicated in classical complement activation by models for C1 complex activation involving shifting of its six Fc-binding C1q subcomponents into a regular hexagonal configuration (**Fig. 2A, 3A**) (12, 13). We therefore hypothesized that 34-1-2S assembles into hexamers on the pulmonary endothelial surface of susceptible mice to trigger complement-dependent acute lung injury.

Imaging methods cannot currently resolve IgG hexamers in vivo, but recent studies have developed methods for inhibiting IgG hexamerization. One approach to impair IgG hexamer assembly is to carbamylate antibodies, converting lysine residues to homocitrullines to alter charge densities in IgG Fc regions, inhibiting Fc:Fc interactions and IgG hexamer assembly (**Fig. 3B**) (23). Mice treated with carbamylated 34-1-2S showed greatly reduced acute lung injury responses compared to littermate controls treated with non-carbamylated 34-1-2S (**Fig. 3C, D**). Carbamylated 34-1-2S was deposited in lungs but, in contrast to unchanged 34-1-2S, did not induce complement C3b/d deposition in the pulmonary microvasculature (**Fig. 3E**)

**Fig. 3.**
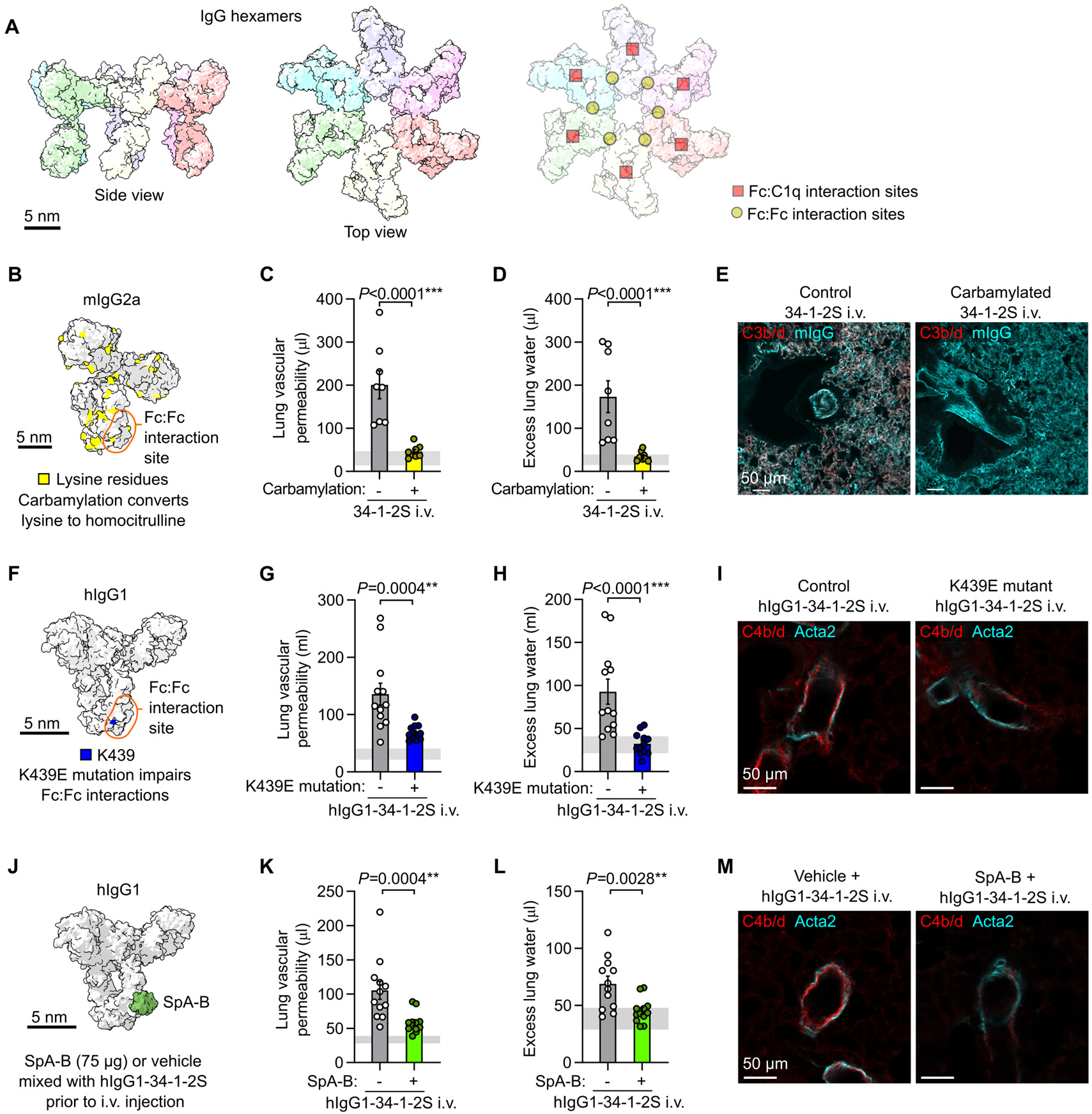
Inhibiting IgG hexamer assembly reduces 34-1-2S-induced acute lung injury responses. **A.** Molecular models of IgG hexamers based on PDB entry 1HZH, showing Fc:Fc and Fc:C1q interaction sites. **B.** Molecular model showing lysine residues in mouse IgG2a (mIgG2a), PDB entry 1IGT. **C.** Lung vascular permeability, **D**. excess lung water measurements and **E**. lung complement C3b/d and mIgG immunostains from LPS-primed BALB/c mice after i.v. injection of carbamylated or control 34-1-2S. **F**. Molecular model showing location of Fc domain lysine 439 (K439) in human IgG1 (hIgG1), PDB entry 1HZH. **G**. Lung vascular permeability, **H**. excess lung water measurements and **I**. lung complement C4b/d and Acta2 immunostains from LPS-primed B6.*H2^d^* mice after i.v. injection with hIgG1-34-1-2S or hIgG1-34-1-2S with K439E mutation. **J**. Molecular model showing binding site of SpA-B to Fc domain of human IgG1 (hIgG1), PDB entries 1HZH and 5U4Y. **K**. Lung vascular permeability, **L**. excess lung water measurements and **M**. lung complement C4b/d and Acta2 immunostains from LPS-primed B6.*H2^d^* mice after i.v. injection with hIgG1-34-1-2S either mixed with 75 µg SpA-B or vehicle control. Samples for injury measurements were collected at 2 hours after antibody injections and lungs were fixed for immunostaining at 5 minutes after antibody injections. Graphs show means ± standard errors. *P*-values are from unpaired two-tailed t-tests on log_10_-transformed data, with group n=8 (**C**, **D**) or n=12 (**G**, **H**, **K**, **L**).

Lysine residues are present on regions of IgG outside of the Fc:Fc interaction interface (illustrated in **Fig. 3B**), so we pursued a more targeted strategy for inhibition of IgG hexamer assembly. We determined the sequence of both heavy and light chain complementary-determining regions and engineered a chimeric antibody with the Fab domain of 34-1-2S fused in frame to human IgG1 (hIgG1-34-1-2S). To test whether hIgG1-34-1-2S causes injury through hexamerization, we also expressed this antibody with an Fc point mutation that inhibits Fc:Fc interactions required for IgG hexamer assembly (K439E, **Fig. 3F**) (15). hIgG1-34-1-2S caused acute lung injury that was reduced by the K439E mutation (**Fig. 3G, H**), as was complement C4b/d deposition in lungs (**Fig 3I**), lending further support to a role for Fc:Fc interactions and hexamerization in the pathogenesis of this disease model.

We also tested a strategy for pharmacologic inhibition of Fc:Fc interactions by mixing hIgG1-34-1-2S with recombinant B domains from *Staphylococcus aureus* protein A (SpA-B), which bind to IgG antibodies near to Fc:Fc interaction sites and inhibit hexamer assembly and complement activation by antibodies targeting bacterial antigens (**Fig. 3J**) (17, 24). We hypothesized that these properties of SpA-B, which likely evolved as part of an immune evasion strategy, might be harnessed to prevent hIgG1-34-1-2S from causing acute lung injury. Adding SpA-B to hIgG1-34-1-2S reduced its ability to both induce acute lung injury (**Fig. 3K, L**) and cause complement C4b/d deposition within pulmonary arterioles (**Fig. 3M**). These findings provide a third line of evidence that Fc:Fc interactions leading to hexamer assembly are important for the injury response caused by this alloantibody.

Turning to hexamer gain of function experiments, the introduction of three mutations into the Fc domain of hIgG1 (RGY mutations: E345R, E430G, S440Y) resulted in antibodies capable of off-target hexamer assembly as well as increased on-target hexamerization (15) (**Fig. 4A**). We hypothesized that RGY-mutated 34-1-2S (RGY-hIgG1-34-1-2S) would have enhanced ability to cause acute lung injury due to increased IgG hexamer formation. We produced RGY-hIgG1-34-1-2S and confirmed its ability to spontaneously assemble into hexamers in solution (**Fig. 4B**). Consistent with a role for alloantibody hexamerization in driving injury, RGY-hIgG1-34-1-2S showed increased potency in triggering acute lung injury relative to hIgG1-34-1-2S (**Fig. 4C, D**), and inducing complement C4b/d deposition in lungs (**Fig. 4E**). A novel chimeric hIgG1 antibody binding a single MHC class I antigen (hIgG1-Kd1, targeting K^d^), did not provoke injury when injected into B6.*H2^d^* mice (**Fig. 4F, G**), consistent with binding to multiple antigens being a requirement for alloantibody-mediated acute lung injury. However, introduction of the RGY mutations promoting hexamerization into this innocuous antibody resulted in a modified version (RGY-hIgG1-Kd1) that was able to provoke increases in lung vascular permeability and edema (**Fig. 4F & G**). Promoting IgG hexamer assembly can therefore increase in vivo pathogenicity of alloantibodies.

**Fig. 4.**
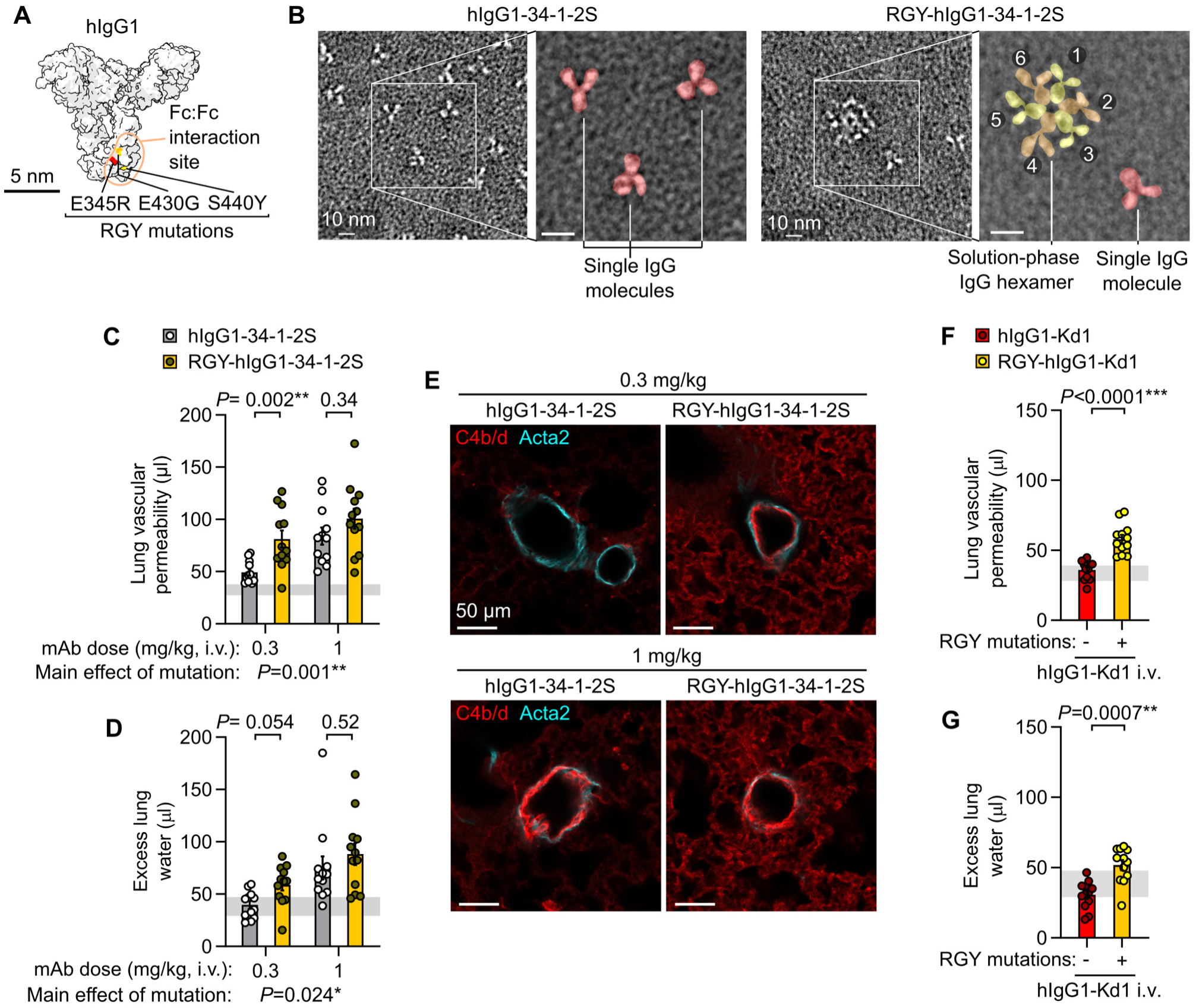
Fc mutations promoting IgG hexamer assembly increase in vivo pathogenicity of alloantibodies. **A.** Molecular model showing amino acids mutated in RGY-hIgG1 antibodies, based on PDB entry 1HZH. **B.** Negative stain electron micrographs showing single hIgG1-34-1-2S molecules and spontaneous solution-phase hexamers formed by RGY-hIgG1-34-1-2S (colored overlay highlights structures in expanded images). **C.** Lung vascular permeability and **D**. excess lung water measurements from LPS-primed B6.*H2^d^* mice injected with control or RGY-mutated hIgG1-34-1-2S monoclonal antibodies (mAbs) at i.v. doses of either 0.3 or 1 mg/kg. **D.** Immunofluorescence staining showing pulmonary arterioles stained for complement C4b/d (red) and Acta2 (cyan) in lung sections from LPS-primed B6.*H2^d^* mice given indicated treatments, with samples fixed 5 minutes after antibody injections. **E.** Lung vascular permeability and **G**. excess lung water measurements from LPS-primed B6.*H2^d^* mice injected with control or RGY-mutated hIgG1-Kd1 (a novel mAb targeting only the H-2K^d^ MHC class I antigen) at 1 mg/kg i.v.. Graphs show means ± standard errors with horizontal line representing standard deviations from ‘no injury’ controls (LPS-primed B6.*H2^d^* mice given hIgG1 isotype control i.v.). Log_10_-transformed data were analyzed using an ordinary two-way ANOVA with Šídák’s multiple comparisons test for effect of Fc mutation within dose level (**C, D**) or unpaired two-tailed t-test (**F, G**), with group n=12.

Another approach for therapeutic exploitation of IgG hexamerization involves use of Fc hexamers as ‘decoy’ treatments. These therapeutic candidates are under investigation as recombinant alternatives to plasma-derived intravenous or subcutaneous immunoglobulin (IVIg or SCIg) treatments that are used in management of autoimmune and alloimmune diseases (25). We hypothesized that due to its ability to inhibit classical complement activation (11, 26), the Fc hexamer ‘decoy’ treatment CSL777 (previously Fc-µTP-L309C) would be effective in preventing alloantibody-mediated acute lung injury.

We randomized mice to receive either CSL777, SCIg (IgPro20, a human plasma-derived immunoglobulin product which is currently used to treat antibody-mediated diseases), or vehicle controls prior to injection with 34-1-2S (**Fig. 5A, B**). Treatment with CSL777 protected mice from developing 34-1-2S-induced lung vascular permeability and pulmonary edema responses at all doses tested, whereas treatment with SCIg only had a partial effect on alloantibody-induced acute lung injury responses (**Fig. 5C-F**). CSL777 treated mice lacked alloantibody-mediated deposition of complement C4 split products on pulmonary arterioles, whereas arteriolar endothelial C4b/d deposition was still observed in lungs of SCIg-treated mice after 34-1-2S antibody injections (**Fig. 5G, H**). Recombinant Fc hexamer therapeutics such as CSL777 might therefore be useful for prevention or treatment of complement-dependent forms of alloantibody-mediated organ injury.

**Fig. 5.**
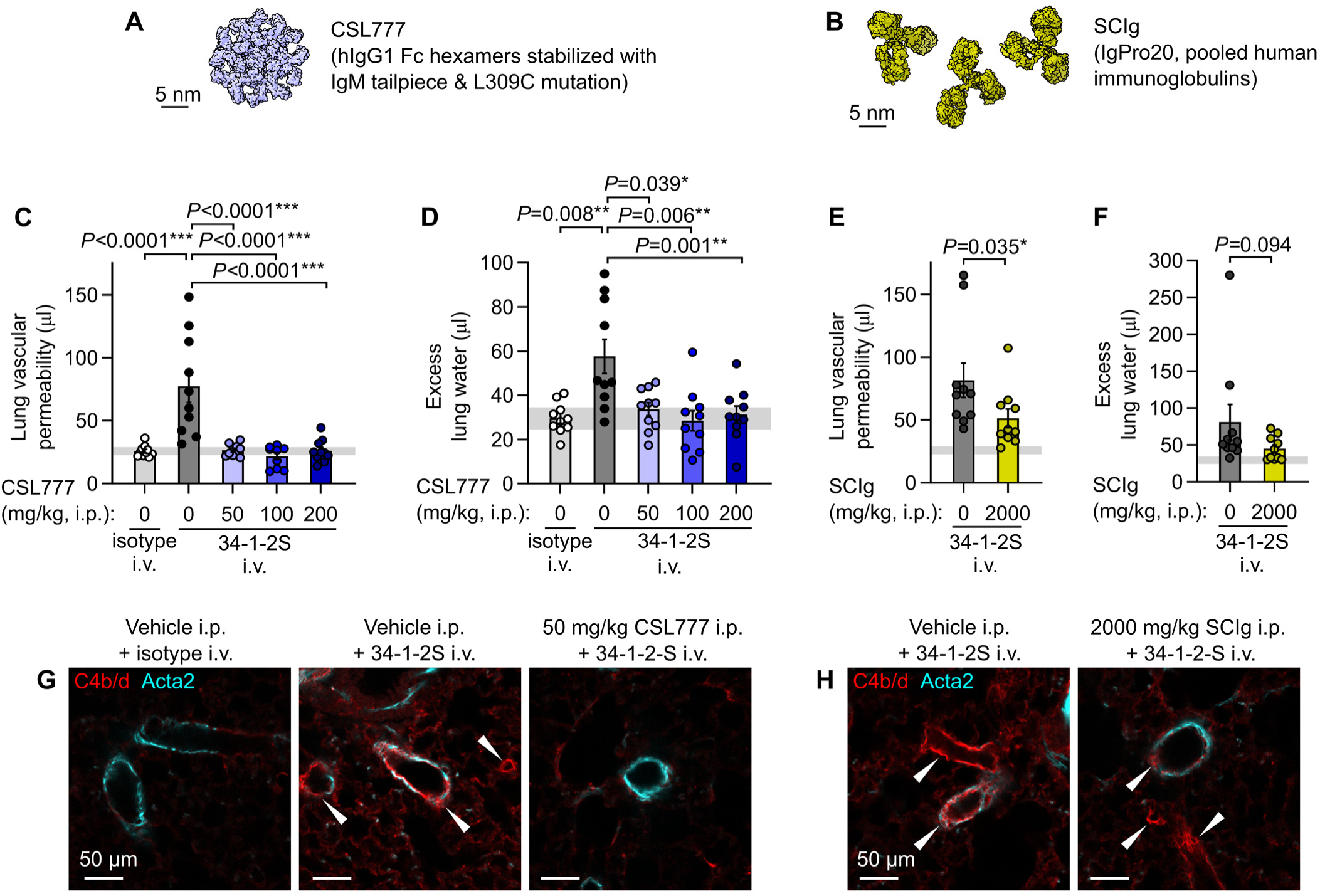
Treatment with recombinant Fc hexamer decoys prevents alloantibody-mediated acute lung injury. **A.** Molecular representation of CSL777, an investigational recombinant Fc hexamer ‘decoy’ treatment which inhibits classical complement activation, based on PDB entry 7X13 (50). **B.** Molecular representation of SCIg, a pooled human immunoglobulin therapeutic with anti-inflammatory properties at high doses, based on PDB entry 1HZH. **C.** Lung vascular permeability and **D**. excess lung water measurements from LPS-primed BALB/c mice given i.p. vehicle or CSL777 at indicated doses 1 hour before i.v. injection with 34-1-2S or mIgG2a isotype control. **D.** Lung vascular permeability and **F**. excess lung water measurements from LPS-primed BALB/c mice given i.p. vehicle or 2000 mg/kg SCIg 1 hour before i.v. injection with 34-1-2S or mIgG2a isotype control. **G**. and **H**. Immunofluorescence showing pulmonary arterioles stained for complement C4b/d (red) and Acta2 (cyan) in lung sections from LPS-primed BALB/c mice given indicated treatments, with samples fixed 5 minutes after antibody injections. White arrowheads point to arterioles with endothelial positivity for C4b/d. Graphs show means ± standard errors with horizontal line representing standard deviations from ‘no injury’ controls (from vehicle + isotype control group). Log_10_-transformed data were analyzed using either (**C**, **D**) an ordinary one-way ANOVA with *P*-values from Dunnett’s multiple comparisons test for difference relative to vehicle + 34-1-2S group, or (**E**, **F**) two-tailed unpaired t-test, with group n=10.

Unlike humans, mice do not express the Fcγ receptor FCGR2A (FcγRIIA, CD32A), negatively impacting the predictive value of mouse models for studying human antibody-mediated diseases (27, 28). To test whether our previous findings held up in a system involving FCGR2A-driven pathology, we crossed existing mouse lines to generate 34-1-2S-mediated injury-susceptible H-2^d^ mice expressing a human FCGR2A (hFCGR2A) transgene (B6.*H2^d^*:*hFCGR2A*^Tg/0^). Mice expressing hFCGR2A developed similar levels of lung injury relative to littermates without hFCGR2A expression but displayed a survival disadvantage (**Fig. 6A-C**), and increased sequestration of platelets in the pulmonary microvasculature (**Fig. 6D,E**). Classical complement activation was still critical for pathogenesis in the presence of hFCGR2A, as knockout of *C1qa* protected mice expressing hFCGR2A from lung injury and mortality (**Fig. 6F-H**). B6.*H2^d^*:*hFCGR2A*^Tg/0^ mice treated with CSL777 were protected from 34-1-2S-mediated injury and showed no mortality responses (**Fig. 6I-K**). These results provide evidence that in a humanized antibody-mediated acute lung injury model involving pathology driven by hFCGR2A, classical complement activation is still a critical event in pathogenesis that can be targeted by therapeutics.

**Fig. 6.**
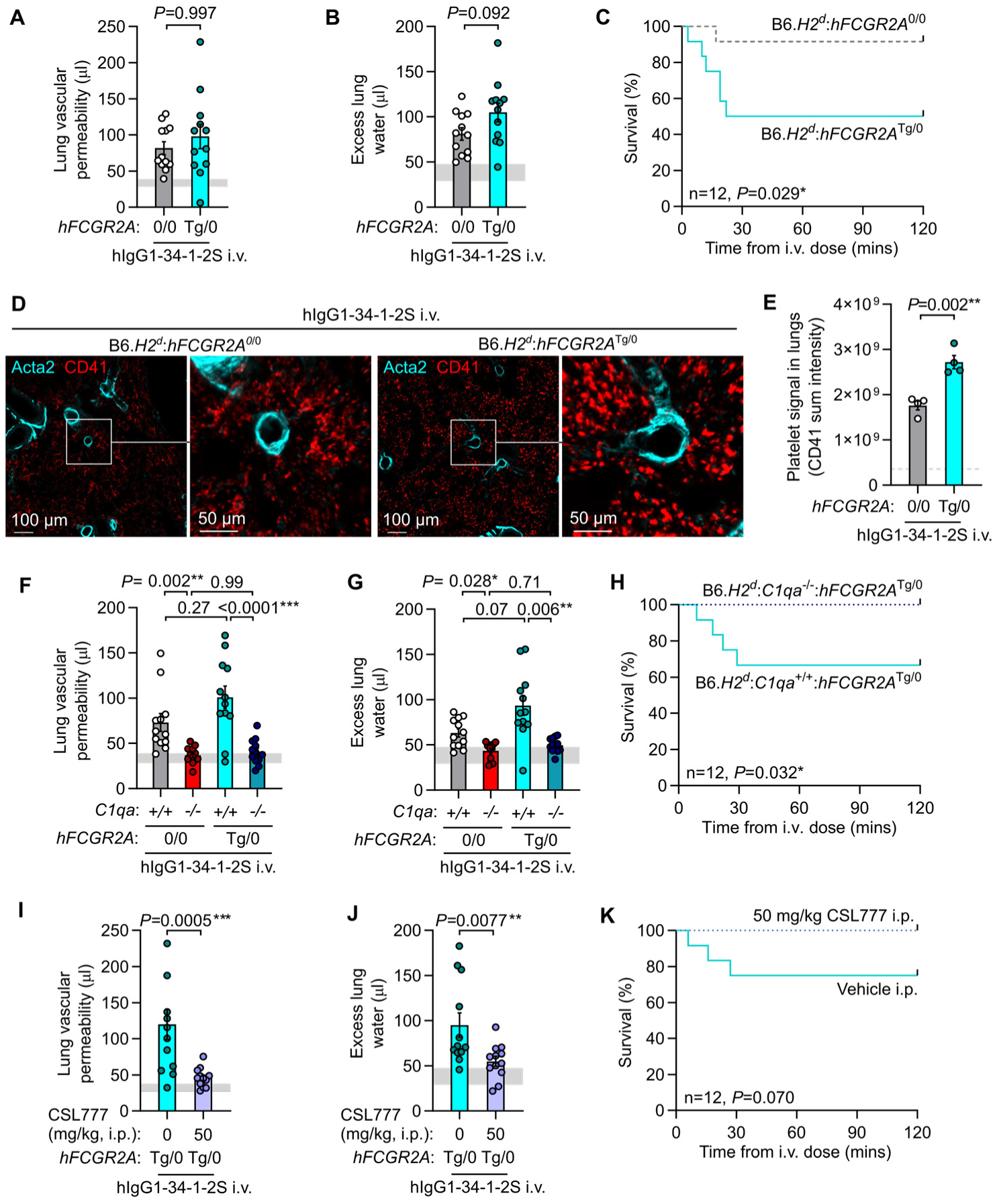
Acute lung injury is complement-dependent in a model incorporating human FCGR2A-mediated pathology. **A.** Lung vascular permeability, **B**. excess lung water and **C**. survival readouts from LPS-primed B6.*H2^d^*:*hFCGR2a*^Tg/0^ mice and B6.*H2^d^* littermate controls given i.v. hIgG1-34-1-2S at 1 mg/kg. **D**. Immunofluorescence imaging of platelet sequestration (CD41, red, with Acta2 in cyan) in lungs from B6.*H2^d^*:*hFCGR2a*^Tg/0^ mice and littermates without hFCGR2A fixed at 20 minutes after hIgG1-34-1-2S injections, quantified in **E**. **F**. Lung vascular permeability, **G**. excess lung water and **H**. survival readouts from LPS-primed B6.*H2^d^*:*C1qa*^+/+^ and B6.*H2^d^*:*C1qa*^-/-^ mice, as well as littermates of each genotype expressing hFCGR2A, given i.v. hIgG1-34-1-2S at 1 mg/kg. **I.** Lung vascular permeability, **J**. excess lung water and **K**. survival readouts from LPS-primed B6.*H2^d^*:*hFCGR2a*^Tg/0^ mice given either vehicle or 50 mg/kg CSL777 i.p. before i.v. hIgG1-34-1-2S at 1 mg/kg. **A, B, E, F, G, I & J** show means ± standard errors with horizontal gray lines showing means or standard deviations of values from ‘no injury’ controls (B6.*H2^d^* mice given LPS i.p. + hIgG1 isotype control i.v.) and were log_10_-transformed prior to analysis. *P*-values are from: (**A, B, I, J**) unpaired, two tailed t-tests; (**F, G**) two-way ANOVA with Šídák’s multiple comparisons test; or (**C, H, K**) log-rank test, with group n=4 (**E**) or 12 (other graphs).

## Discussion

This study advances our understanding of immunology in three areas. Our work elucidates the molecular determinants of susceptibility in a widely used inflammation model. To our knowledge, our experiments represent the first in vivo evidence that alloantibody hexamerization is important for pathophysiology. In addition, we show that two experimental therapeutic approaches that target antibody hexamerization can prevent alloantibody-mediated organ injury.

The findings presented in this study allow us to explain a long-standing mystery of great mechanistic importance in a widely used model of immune-mediated organ injury; in particular, why 34-1-2S antibody injections (but not injections of other anti-MHC class I antibodies) cause such striking pathophysiology in mice carrying the H-2^d^ but not H-2^b^ MHC haplotype (6, 19, 20, 28–38). We posit that high affinity binding to multiple MHC class I antigens on the pulmonary endothelium of mice with the H-2^d^ haplotype facilitates sufficiently dense alloantibody deposition for IgG hexamer assembly to occur.

These IgG hexamers then direct classical complement activation onto the pulmonary endothelial surface, initiating the excessive leukocyte and platelet responses that have been reported in previous studies to cause acute lung injury in this model (6, 19, 20). Furthermore, we were able to render an innocuous antibody specific for a single MHC class I antigen into a pathogenic antibody by introducing mutations promoting increased on and off-target hexamerization. An additional implication of our findings is that both lymphocyte crossmatch and single antigen bead assays for detecting complement-fixing antibodies may lack sensitivity for detecting antibodies that activate complement in vivo. The mobility, density and diversity of antigens on the lymphocytes or solid-phase beads used in these assays does not exactly resemble those on endothelial cell surfaces targeted by donor-specific antibodies in vivo, and our results indicate that each of these factors determines complement-fixing capability of antibodies. Conversely, antigen density on beads exceeding that found in vivo may give false positive findings of complement-fixing antibody responses.

By demonstrating that IgG hexamers are important in pathophysiology and represent therapeutic targets in vivo, our work builds on in vitro studies implicating antibody hexamerization in complement activation by antibodies targeting antigens on liposomes, lymphoma cells or bacterial membranes (15–18, 23, 39). Further clinical translation will require determination of the importance of IgG hexamers in more complex models of diseases involving complement-activating alloantibodies (e.g. AbMR, TRALI, hemolytic transfusion reactions, and hemolytic disease of the fetus and newborn) or autoantibodies (e.g. warm autoimmune hemolytic anemia, antiphospholipid syndrome, myasthenia gravis and neuromyelitis optica). As SpA-B does not inhibit IgG3-mediated complement activation but IgG3 can assemble into hexamers (15, 16), it will also be important to develop strategies to inhibit IgG3 hexamerization to examine the therapeutic potential of targeting hexamers formed by IgG3 antibodies.

Our results also provide new insights into the modes of action of past, present, and potential future therapeutics. Full length staphylococcal protein A (SpA) has been used as a therapeutic in the form of a now-discontinued extracorporeal immunoadsorption product (Prosorba column). Efficacy of SpA immunoadsorption has been observed in patients with symptoms unchanged by plasma exchange, an effect ascribed to leakage of SpA from columns into the bloodstreams of patients resulting in B cell depletion caused by the action of SpA as a B cell receptor super agonist (40). Purified SpA infusions (PRTX-100) were subsequently studied in early-stage clinical trials before abandonment for financial reasons (41). Our results suggest that there may be settings where therapeutics based on the SpA-B subdomain of SpA have efficacy through preventing complement activation without risk of adverse effects related to immune complex formation and B cell super agonism caused by immunoglobulin polyvalency of full-length SpA. Donor-derived immunoglobulin products (e.g. IVIg and SCIg) are currently used to treat antibody-mediated disease flares. Our observation that SCIg reduces injury responses but does not prevent classical complement activation in vivo is concordant with previous studies concluding that immunoglobulin therapeutics act on downstream mediators in vitro and in vivo (33, 42). CSL777 is an attractive potential future therapeutic for treatment of alloantibody-mediated diseases as it showed efficacy in our models, and had a mode of action that would be anticipated to prevent complement activation by both IgM and IgG antibodies as well as pathophysiology resulting from Fcγ receptors (11, 25, 26). CSL777 also lacks issues with use of donor-derived products related to sourcing, purification and concentration for injections (25, 43).

In conclusion, this study provides evidence that IgG hexamers can be important triggers of complement-dependent pathophysiology in vivo. Our preclinical studies support further investigation of IgG hexamerization inhibitors and IgG hexamer ‘decoy’ therapeutics for use in preventing disease states caused by antibodies and complement activation.

## Materials and methods

### Animals

B6.C-*H2^d^*/bByJ (B6.*H2^d^*) mice (Cat# 000359) (44) and B6(Cg)-*C1qa^tm1d(EUCOMM)Wtsi^*/TennJ (*C1qa*^-/-^) mice (Cat# 031675) (22) originated from the Jackson Laboratory. B6-background mice were bred with B6.*H2^d^*mice and progeny were crossed to produce mice with homozygous expression of H2^d^ MHC antigens for use in experiments (6). B6.ConK^d^-on and B6.Tg(CD2-Tcra,-Tcrb)75Bucy (TCR75) mice were provided by J. Zimring (21, 45). BALB/c mice were from Charles River Laboratories (Cat# 028). B6;SJL-Tg(FCGR2A)11Mkz/J mice (expressing human FCGR2A isoform R131, Jackson Laboratory Cat# 003542) (27) were backcrossed to B6 congenicity (46). Male mice were used as female mice are not susceptible to 34-1-2S-mediated injury (5). Mice were studied at 8-16 weeks of age after maintenance in the UCSF specific pathogen-free animal facility for at least 2 weeks. Procedures received ethical approval from the UCSF IACUC committee.

### Surface plasmon resonance

Binding affinities were determined by injecting serial dilutions (0.5-200 nM) of MHC class I monomers (MBL International Cat#: TB-5001-M (K^b^ presenting SIINFEKL); TB-5008-M (D^b^ presenting RAHYNIVTF); TB-M552-M (K^d^ presenting VYLKTNVFL); TB-M536-M (D^d^ presenting IGPGRAFYA); TB-M521-M (L^d^ presenting SPSYVYHQF)) over monoclonal antibodies bound via amine coupling to SensEye G Easy2Spot sensors (Ssens Cat# 1-09-04-006), assayed in triplicate with an IBIS MX96 SPR imager.

### Alloantibody-mediated acute lung injury model

As previously described, mice were given i.p. injections of LPS (Sigma Aldrich Cat# L2880, 0.1 mg/kg) for ‘priming’ needed to render barrier-housed mice responsive to antibody injections (19). At 24 hours after LPS priming, mice were anesthetized (0.6 mg/kg ketamine + 0.1 mg/kg xylazine i.p.) and the indicated antibody treatments were injected into the jugular vein over the course of 1 minute (at 1 mg/kg unless otherwise stated). Antibodies were from BioXCell (mIgG2a isotype control Cat# BE0085, 34-1-2S Cat# BE0180, hIgG1 isotype control Cat# BE0297) or newly produced (described below). Lung vascular permeability was measured by giving each mouse 0.01 KBq of ^131^I-conjugated albumin (Iso-Tex Diagnostics, NDC:50914-7731) together with i.v. antibody injections, collecting lungs and blood samples 2 hours later or at cessation of breathing for radioactivity measurements used to quantity volume of extravasated plasma in lungs (lung vascular permeability). Wet-dry weight ratios of lungs and blood were used to calculate excess lung water volumes (6).

### Immunofluorescence imaging

Cryosections were made at 200 µm or 400 µm thickness from lungs fixed by inflation with and immersion in 1% formaldehyde in PBS, as previously described (6). Sections were incubated overnight with antibodies targeting C3b/d (Novus Cat# NB200-540), C1qa (Abcam Cat# ab182451), C4b/d (Novus Cat# NB200-541), Scgb1a1 (Sigma Aldrich Cat# ABS1673) or CD41 (Biolegend Cat# 133939) together with a FITC-conjugated antibody raised against Acta2 (Sigma-Aldrich Cat# F3777) were incubated overnight at 1:500 with 5% normal donkey serum, 0.1% bovine serum albumin and 0.3% triton X-100 in phosphate-buffered saline (PBS). After washing, Cy3 or Alexa Fluor 647-conjugated cross-adsorbed polyclonal secondary antibodies targeting rat, rabbit, goat and/or mouse IgG (Jackson Immunoresearch Cat# 712-165-153, Cat# 711-165-152, Cat# 705-605-147 and/or Cat# 115-605-206) were incubated with sections at 1:500 in PBS + 0.3% triton X-100 overnight. After additional washes, sections were either mounted in Vectashield (Vector Laboratories Cat# H-1700) for standard confocal imaging on a Nikon A1r microscope, or cleared after staining using the EZ clear protocol (47) for 3D imaging with a Nikon AZ100M confocal microscope.

### Antibody carbamylation

Carbamylation of 34-1-2S was achieved by incubating 1 mg of antibody in PBS + 0.1 M KOCN for 1 hour at 37°C before buffer exchange back into PBS (23). Control 34-1-2S was subjected to the same process with omission of KOCN.

### Antibody sequencing and engineering

The 34-1-2S hybridoma was purchased from ATCC (Cat# HB-79). To generate the Kd1 hybridoma, B6 mice were injected i.v. with 3×10^6^ CD4+ T cells from TCR75 mice transgenic for a T cell receptor (TCR) specific for a peptide from K^d^ presented by IA^b^, and i.p. with 5×10^6^ Con-K^d^-on splenocytes, resulting in an extreme B cell antibody response directed at an immunodominant peptide from K^d^, the only mismatched antigen between donor and recipient. Three days after a boost with an additional i.p. injection with Con-K^d^-on splenocytes, splenocytes from the sensitized recipient were fused with a myeloma cell line as previously described (48), and monoclonal antibodies specific for K^d^ were identified using Con-K^d^-on splenocytes as targets (21).

Monoclonal antibody aliquots were digested with either peptidyl-Asp metalloendopeptidase, chymotrypsin, elastase, trypsin, or pepsin enzymes. Peptides were then assayed using liquid chromatography coupled to tandem mass spectrometry for sequencing of variable fragments (Bioinformatics Solutions Inc.). Amino acid sequences determined were, for 34-1-2S:

EVQLQQSGAEFVRPGASVKLSCTASGFNLKDDYMFWVKQRPEQGLEWIGWIAPDNGDTEYASKFQGKATITADTSSNTAYVQLSSLTSEDTAVYYCTTWGYYSYVNYWGQGTTLTVSS (heavy chain variable region) and: DIQMTQSPSSLSASLGERVSLTCRASQDIGSNLNWLQQEPDGTIKRLIYATYSLDSGVPKRFSGSRSGSDYSLTISSLESEDFVDYYCLQYASSPYTFGGGTKLEIK (light chain variable region); and for Kd1: EVLLVESGGDLVKPGGSLKLSCAASGFTFRTYAMSWVRQTPEKRLEWVATIGDDGSYTFYPDNVKGRFTISRDNAKNNLYLQMRHLKSEDTAIYFCARDGLFAYWGQGTLVTVSA (heavy chain variable region) and:

DIQMTQSPSSLSASLGGKVTITCKASQDIKKNIAWYQYKPGKGPRLLIHYTSTLQPGISSRFSGSGSGRDYSFSISNLEPEDIATYYCLQYDSLLYTFGGGTKLEIK (light chain variable region). Correct identification of variable domains was confirmed by performing sequencing of the products of 5’ rapid amplification of cDNA ends (5’-RACE) for heavy and light chain mRNA using RNA isolated from hybridomas. The isolated 5’-RACE amplicons contained open reading frames that encoded the above peptides sequenced by mass spectrometry. These sequences were codon-optimized and antibodies were expressed as chimeric hIgG1 with or without Fc point mutations in a HEK293 cell system and purified using protein A and buffer exchange (Absolute Antibody).

### Pharmacologic treatments

Recombinant subdomain B from Staphylococcus aureus (SpA-B, amino acid sequence: HHHHHHADNKFNKEQQNAFYEILHLPNLNEEQRNGFIQSLKDDPSQSANLLAEAKKLNDAQAPK, His-tag added for purification) was produced in an *E. coli* system by Genscript and supplied in protein storage buffer (50 mM Tris-HCl, 150 mM NaCl, 10% Glycerol, pH 8.0). At 1-2 hours before i.v. injection, SpA-B (3 mg/kg) or vehicle were mixed with hIgG1-34-1-2S resulting in a 30% vehicle, 70% PBS mixture.

Trial formulations of CSL777 (in PBS vehicle), as well as clinical-grade IgPro20 (Hizentra™) and the proprietary vehicle for IgPro20 were provided by CSL Behring. Mice were given i.p. injections of CSL777, IgPro20 or relevant vehicle 1-2 hours before i.v. injections of antibodies at stated doses.

### Negative stain electron microscopy

Antibody samples were diluted to 0.01 mg/ml in 25 mM HEPES, 100 mM NaCl and added to carbon-coated grids (TedPella Cat# 01702-F, manually coated with 20 nm carbon using a Cressington 208 Sputter Coater). Sample-coated grids were stained using 0.75% uranyl formate and imaged on an FEI Tecnai T12 transmission electron microscope.

### Experimental design and analysis

Within-cage randomization was used for group allocations in studies testing exogenous treatments. Littermate controls from heterozygous crosses were used to test the effect of *C1qa* knockout. Congenic animals housed in the same room were used to study haplotype effects. Handlers were blinded to group allocations during experiments. Group numbers (n) and analysis approaches were predetermined before initiation of experiments. Statistical tests used on each dataset are described in figure legends.

### Software

GraphPad Prism was used for graphing and statistical analysis. UCSF ChimeraX was used for molecular graphics (49). Imaris was used to render fluorescence micrographs and ImageJ was used to process electron microscopy data.

## Acknowledgements

Imaging studies were possible thanks to the UCSF Biological Imaging Development Colab, and UCSF Electron Microscopy core. Sandro Prato (CSL Innovation Pty Ltd, Parkville, Victoria, Australia) provided advice on in vivo administration of CSL777 and SCIg.

## Funding

National Institutes of Health grant R01 HL138673 (JCZ and MRL)

National Institutes of Health grant R35 HL161241 (MRL)

Association for Advancement of Blood and Biotherapies (AABB) Foundation (formerly National Blood Foundation) Early Career Scientific Research Grant (SJC)

Canadian Institutes of Health Research (ÉB)

## Authorship

Conceptualization: SJC, JCZ, MRL

Methodology: SJC, MRL, AEHB, GV, ÉB, RS, JCZ

Investigation: SJC, YS, JJT, NK, DPB, AEHB

Funding acquisition: SJC, JCZ, MRL

Writing – original draft: SJC, MRL

Writing – review & editing: SJC, YS, JJT, NK, DPB, AEHB, GV, ÉB, RS, JCZ, MRL

Conflict of interest statement: RS is an employee of CSL Behring AG, Switzerland. The authors have no additional conflicts of interest.

## Data and materials availability

All data are available in the main text or the supplementary materials.

## Supplementary materials

Supplementary materials and methods.

Supplementary Figures:

**Fig. S1. Binding of MHC class I monoclonal antibodies to MHC class I monomers.**

**Movie S1. Complement C4 split product deposition in the pulmonary vasculature.**

