## Supplemental materials and methods, Fig. S1, Movie S1 legend for "IgG hexamers initiate acute lung injury"

### Supplementary materials and methods

#### *Additional MHC class I antibodies*

Additional IgG2a monoclonal antibodies targeting MHC class I antigens used in surface plasmon resonance (SPR) experiments were: clone AF6-88.5.5.3 (BioXCell Cat# BE0121); clone 20-8-4S, produced locally from hybridoma (ATCC Cat# HB-11); clone SF1.1.10 (BioXCell Cat# BE0104); clone 30-5-7S (Thermo Fisher Cat# MA1-70109); clone 34-5-8S (Thermo Fisher Cat# MA1-70108).

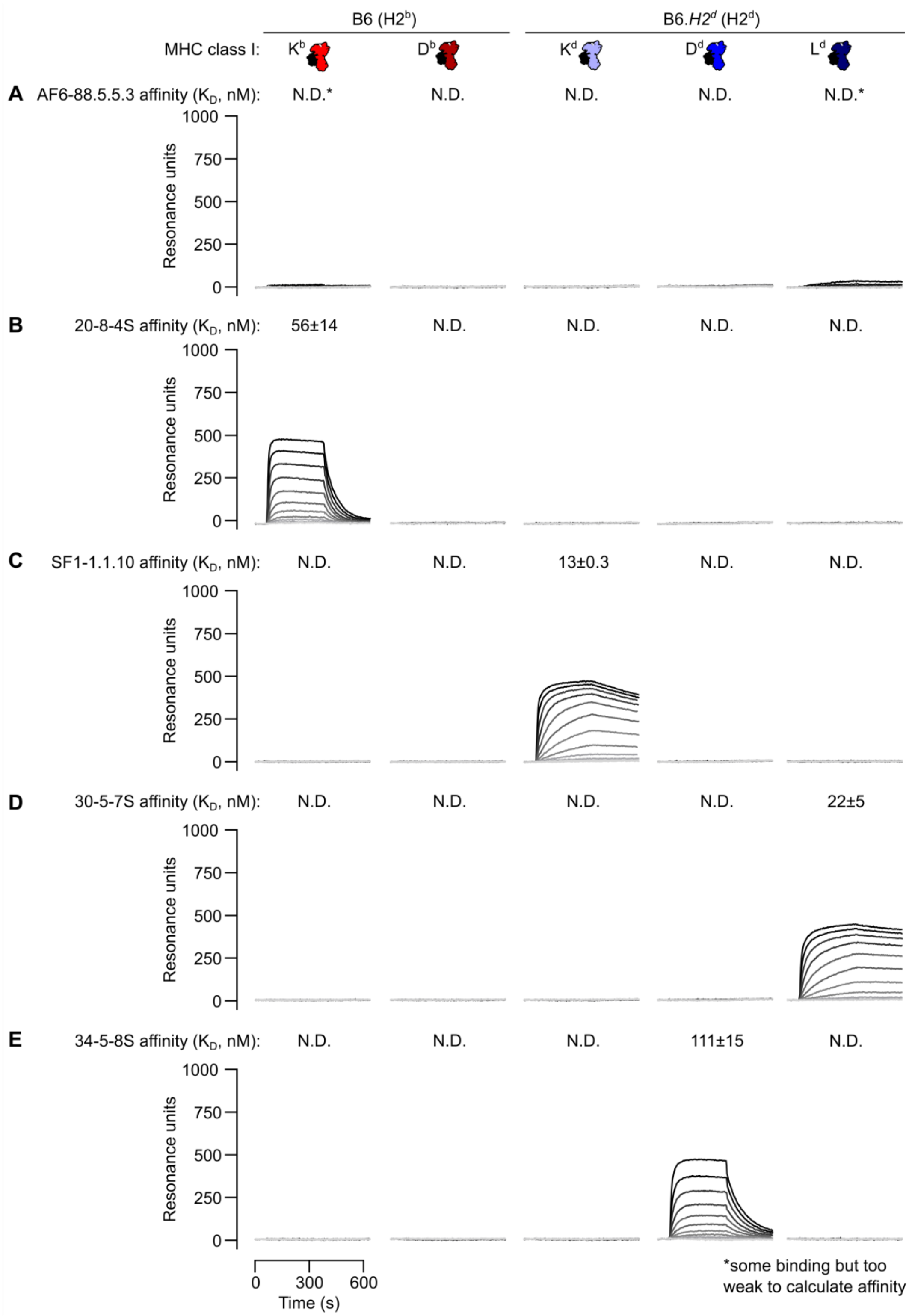


**Fig. S1. Binding of MHC class I monoclonal antibodies to MHC class I monomers.**

Representative surface plasmon resonance (SPR) sensorgrams showing binding of antibody clones: (A) AF6-88.5.5.3; (B) 20-8-4S; (C) SF1-1.1.10; (D) 30-5-7S; and (E) 34-5-8S to each of the classical MHC class I antigens expressed by mice with $H2^b$ or $H2^d$ haplotypes. Dissociation constants ($K_D$) are given as means ± standard deviations (n=3).

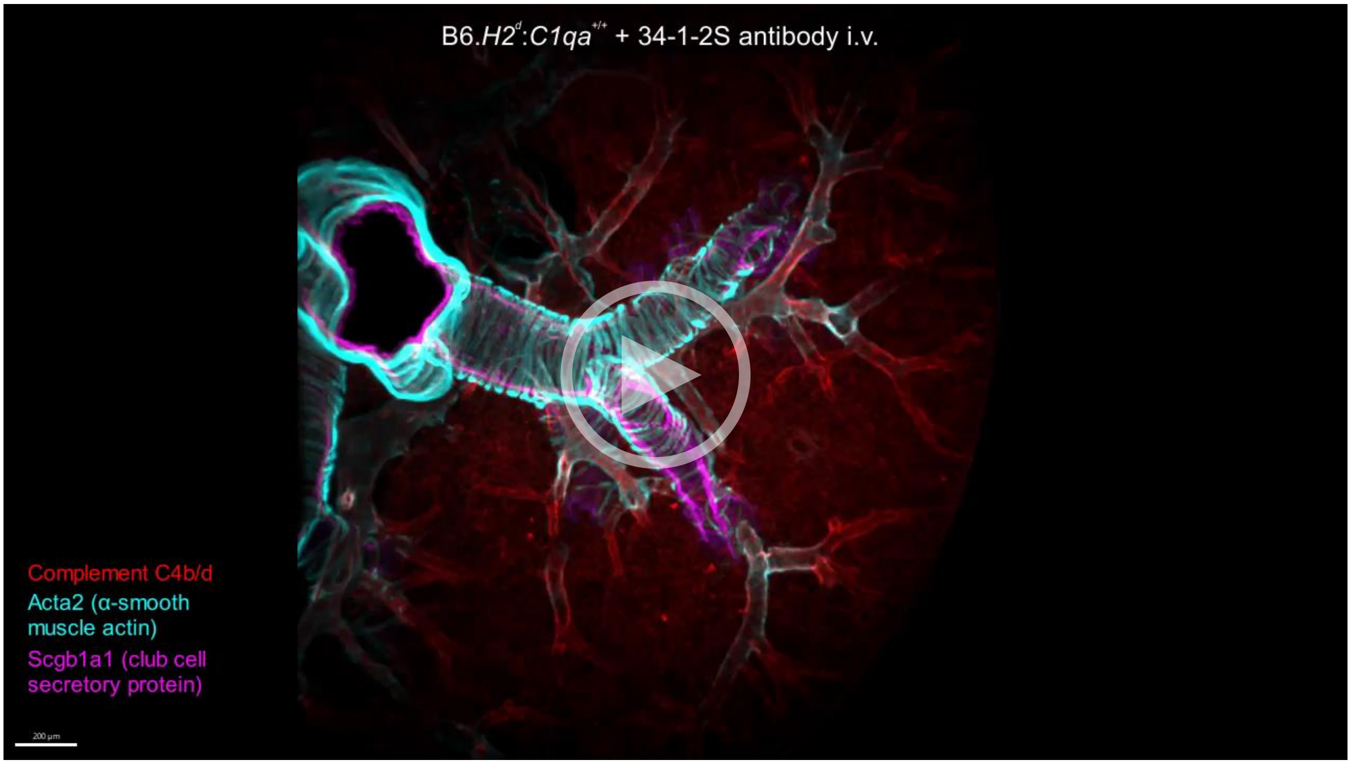


**Movie S1. Alloantibody-mediated complement C4 split product deposition in the pulmonary vasculature.** LPS-primed B6.*H2*[d] mice of either *C1qa*$^{+/+}$ or *C1qa*$^{-/-}$ genotype were given 34-1-2S or its isotype control i.v. at 1 mg/kg. At 5 minutes after antibody injections, lungs were collected, fixed, sectioned at 400 µm thickness, immunostained and cleared using the EZ clear protocol. Images show staining for complement C4b/d (red), Acta2 (α-smooth muscle actin surrounding airways and resistance arterioles, cyan) and in some samples, Scgb1a1 (club cell secretory protein, abundant on epithelium of medium-sized airways, magenta). Detail expands highlight arterioles that show strong endothelial positivity for complement C4b/d in C1qa-sufficient mice injected with 34-1-2S antibody. Movie file attached as supplementary data.
